# Psilocybin acutely dissociates cognitive and affective empathy, with empathic concern predicting persisting positive psychological effects

**DOI:** 10.64898/2026.08.04.742704

**Authors:** N.L Mason, P Mallaroni, K.P.C Kuypers, R De La Torre Fornell, J.T Reckweg, K.H Preller, J.G Ramaekers

## Abstract

**Background:** Psychedelics have been proposed to enhance empathy and social connectedness, but it remains unclear whether these effects reflect global increases in empathic ability, whether they persist beyond the acute drug state, and whether they contribute to later psychological outcomes.

**Methods:** In a randomized, double-blind, placebo-controlled, parallel-group study, healthy participants received psilocybin (0.17 mg/kg) or placebo. Three dissociable components of empathy were assessed using the Multifaceted Empathy Test at baseline, during the acute drug state, and 7 days later: cognitive empathy (accuracy of emotion identification), emotional arousal (affective activation in response to another’s emotional state) and empathic concern (other-oriented compassion toward the depicted person. Persisting positive psychological effects were assessed at follow-up. Circulating oxytocin was measured, and spectral dynamic causal modeling was applied to resting-state fMRI data acquired during the acute drug state to examine effective connectivity within an empathy-relevant network.

**Results:** Compared with placebo, psilocybin acutely reduced cognitive empathy selectively for positive stimuli, while increasing emotional arousal for positive stimuli and empathic concern across valences. These empathy-related effects did not persist 7 days later. However, acute increases in empathic concern mediated later positive psychological changes, including improved attitudes about life and self, mood, and relationships. Psilocybin also increased circulating oxytocin compared to baseline, but oxytocin changes were not associated with empathy changes. Effective-connectivity analyses showed that empathic responding under psilocybin was associated with altered directional coupling among superior temporal and parahippocampal regions.

**Conclusions:** Psilocybin does not simply enhance empathy, rather it impairs the identification of positive emotional states while heightening affective engagement and concern. Although these effects do not persist, acute empathic concern may contribute to later positive psychological change.

## Introduction

Impairments in social cognition and empathic responding are common across psychiatric disorders and are increasingly recognized as important contributors to interpersonal dysfunction, social withdrawal, and reduced treatment response (1–3). Empathy is not a unitary construct but comprises partially dissociable cognitive and affective components. Cognitive empathy refers to the capacity to explicitly infer or identify another person’s emotional state, overlapping conceptually with mentalizing or theory of mind. At the affective level, emotional empathy refers to affective responses elicited by another person’s emotional state, including emotional resonance and arousal (4–9). Closely related, but conceptually distinct, is empathic concern, which reflects other-oriented feelings of care, warmth, or compassion toward another person(7). These capacities support prosocial behavior, interpersonal cooperation, and adaptive social functioning (4, 10), whereas disruptions in social-cognitive and affective interpersonal processes are observed across multiple psychiatric conditions (1, 2). Identifying interventions that can alter distinct components of empathic responding may therefore be relevant for understanding, and potentially improving, social functioning in mental health.

Psychedelic compounds have been proposed to acutely alter social and affective processing. Across studies, classic serotonergic psychedelics such as psilocybin, lysergic acid diethylamide (LSD), and ayahuasca, as well as related compounds such as 3,4-methylenedioxymethamphetamine (MDMA), have been associated with acute changes in empathy and related social processes (for a review see Preller and Vollenweider (11)). However, these effects do not appear to reflect a global enhancement of empathy but rather selectively alter its subdomains. Empathic concern and emotional arousal are more consistently increased following psychedelic administration (12–19), whereas findings on cognitive empathy have been more mixed and may depend on stimulus valence (12, 13, 18, 20, 21). These findings suggest that psychedelics may shift the balance of empathic responding toward affective engagement with others’ emotions and away from explicit social-cognitive inference, rather than uniformly improving empathic ability. An important unresolved question is whether psychedelic-induced changes in empathy extend beyond the acute drug state and whether they contribute to lasting psychological wellbeing. Acute empathy changes could be relevant in at least two ways. First, they may persist directly after drug effects have subsided, producing sustained changes in empathic functioning. Second, even if they do not persist as stable trait-like changes, acute shifts in empathy may catalyze broader psychological or interpersonal processes that unfold over subsequent days. Evidence for persisting empathy-related effects remains limited. Naturalistic studies have reported increases in affective empathy several days after psilocybin or ayahuasca use in ceremonial or retreat settings (22, 23), and recent clinical work suggests that psilocybin-assisted therapy may increase empathic concern in patients with major depression(24). However, these studies involved highly social or therapeutic contexts, making it difficult to determine whether persisting effects generalize to controlled experimental settings without structured interpersonal intervention. Moreover, it remains unclear whether acute drug-induced changes in empathy are related to the positive psychological changes reported after psychedelic experiences (for a review see Aday, Mitzkovitz (25)).

The biological mechanisms underlying psychedelic-induced changes in empathy are also poorly understood. Cognitive empathy, emotional empathy, and empathic concern rely on partially overlapping but dissociable neural systems involving regions implicated in social perception, mentalising, autobiographical, affective or contextual processing(4, 5). These include superior temporal, paracingulate, and medial temporal regions that may interact differently depending on whether empathic responding requires explicit emotion identification, affective resonance, or other-oriented concern. Because psychedelics acutely alter large-scale brain organization and the balance between higher-order associative and sensory-affective systems(26), they may provide a useful pharmacological model for dissociating cognitive and affective components of empathy at the neural level. Dynamic causal modeling is well suited to this question, as it estimates directed rather than undirected connectivity(27). This approach may therefore clarify whether psilocybin-induced changes in empathic responding are associated with altered information flow within an empathy-relevant network. In parallel, oxytocin has been implicated in social-affective behavior and empathic responding(28), and psychedelic administration has been shown to increase circulating oxytocin concentrations(29). However, whether peripheral oxytocin changes explain psychedelic-induced alterations in empathy remains uncertain(15, 29).

The present study investigated the acute and persisting effects of psilocybin on dissociable components of empathic responding in a randomized, double-blind, placebo-controlled, parallel-group design. Empathy was assessed using the Multifaceted Empathy Test at baseline, during the acute drug state, and 7 days later, allowing separate assessment of cognitive empathy, emotional arousal, and empathic concern(30). We first tested whether psilocybin altered these empathy components acutely and whether any changes persisted at follow-up, compared to baseline and to the placebo condition. We then examined whether acute changes in empathy statistically mediated the relationship between psilocybin administration and persisting positive psychological effects. Finally, we assessed candidate biological mechanisms associated with acute empathy changes by measuring circulating oxytocin concentrations and applying spectral dynamic causal modeling to resting-state functional MRI data acquired during the acute drug state.

## Materials and Methods

A detailed description of the experimental procedure is provided in the Supplementary Methods and briefly summarized here.

The present study was conducted between July 2017 and June 2018 at Maastricht University, employing a balanced randomized (1:1), placebo-controlled, double-blind, parallel-group design. Sixty healthy participants, with previous experience with a psychedelic drug but not within the past 3 months, were allocated to a treatment condition (0.17 mg/kg psilocybin or placebo, p.o.). Groups were matched for age, sex, and education level.

Participants visited the lab on three separate occasions. The first visit included familiarizing with testing day procedures and completing the empathy task (baseline), and satisfaction with life questionnaire. The second visit consisted of the formal testing day, with treatment administration (psilocybin or placebo), fMRI, sampling of psilocin and oxytocin concentrations, assessment of satisfaction with life, and completion of the empathy task (acute). The third testing day took place 7 days after the acute testing day and included the final completion of the empathy task (follow-up) and the completion of the persisting effects questionnaire.

This study was conducted according to the code of ethics on human experimentation established by the declaration of Helsinki (1964) and amended in Fortaleza (Brazil, October 2013) and in accordance with the Medical Research Involving Human Subjects Act (WMO) and was approved by the Academic Hospital and University’s Medical Ethics committee. All participants were fully informed of all procedures, possible adverse reactions, legal rights and responsibilities, expected benefits, and their right for voluntary termination without consequences. The present data were part of a larger clinical trial (Onderzoekmetmensen; https://onderzoekmetmensen.nl/nl/trial/25919; NL-OMON25919; 22-03-2017), of which parts have been previously published (31–33).

### Multifaceted Empathy Test

The multifaceted empathy test (MET) consists of 40 pictures of people in various emotional states, with 50% being positive and 50% negative (34). It assesses three dissociable components of empathic responding. Cognitive empathy (CE) was assessed by asking participants to select, from four options, the emotion word which matched the depicted emotion. Emotional arousal, operationalized as *implicit emotional empathy* in the MET, was assessed by asking participants to rate on a scale from 1 to 9 “how aroused does this picture make you feel”. Empathic concern, operationalized as *explicit emotional empathy* on the MET, was assessed by asking participants to rate on the same scale “how concerned do you feel for this person”. The number of correctly classified pictures (CE) and the concern and arousal ratings per valence and averaged across valences were used as dependent variables. Previous validity and reliability analysis of the MET have shown to be in the good to highly satisfactory range (34), and previous studies have found it to be sensitive to the effects of psychedelics (12, 14–18, 22).

### Satisfaction with Life Scale

The Satisfaction with Life Scale (SWLS) is a five-item questionnaire designed to measure global cognitive judgments of satisfaction with one’s life (35). It has been used to measure the life satisfaction component of subjective well-being (35, 36).

### Persisting Effects Questionnaire

The Persisting Effects Questionnaire (PEQ) is a 143-item long scale developed to assess changes in attitudes, moods, behavior, and spiritual experience (37, 38) following a psychedelic experience. Prior research has found that the PEQ is sensitive to the prolonged effects of psychedelics in healthy individuals, occurring even one year after ingestion (39).

### Oxytocin

Venous blood samples were taken at baseline, and approximately 80 and 150 minutes post-treatment to assess oxytocin concentrations. See Supplemetary Materials for details.

#### Neuroimaging acquisition and preprocessing

Resting-state 7T fMRI data acquired during the acute drug state (102 min post) were used to assess whether psilocybin altered directed interactions within an empathy-relevant network (see Supplementary for acquisition details).

Five regions of interest were selected to capture directed interactions among social-perceptual, mentalizing, and contextual-memory systems relevant to empathic responding. Regions were defined as 8-mm spheres centered on Neurosynth-derived MNI peak coordinates: paracingulate cortex (ParaCing: +2, +28, +22), left superior temporal gyrus (lSTG: −58, −32, +8), right superior temporal gyrus (rSTG: +56, −12, 0), left parahippocampal cortex (lParaHip: −26, −38, −12), and right parahippocampal cortex (rParaHip: +24, −40, −12). Paracingulate and superior temporal regions are consistently implicated in theory of mind, social perception, and interpretation of others’ actions, intentions, and emotional states(40, 41), with superior temporal activity further differentiating cognitive and emotional empathy during the Multifaceted Empathy Test(42). The parahippocampus was included to represent the contextual-associative and self-referential memory processes on which empathic judgements draw(43), processes also implicated in psychedelic-induced changes in self-related processing and connectedness(31, 44, 45).

Regional time series were extracted and denoised (motion, CSF, and white-matter regressors; 128-s high-pass filter; no global signal regression), with no residual motion differences between groups after excluding participants with mean framewise displacement > 0.5 mm (see Supplementary Methods)

#### Spectral dynamic causal modelling (DCM)

Spectral DCM fits a biophysical state-space model to the observed cross-spectra of BOLD signals, to estimate underlying neuronal states and the rate of change in neural activity in each region (in hertz) as a function of activity in other regions(46). To estimate directed interactions, a fully connected spectral DCM (SPM12) with no exogenous inputs was specified per subject and fitted to the cross-spectral density of the BOLD signal, yielding directed coupling estimates between regions and each region’s intrinsic (log-scaled, inhibitory) self-connection. Participants with model explained variance below 60% were excluded, resulting in a final DCM sample of 52 participants (psilocybin: n = 23; placebo: n = 29). Model fit did not differ between treatment groups (explained variance: psilocybin: 85.66 ± 6.27%; placebo: 86.51 ± 4.00%; Welch’s t(35.6) = −0.57, p = .58; Wilcoxon p = .93).

#### Parametric Empirical Bayes (PEB)

Subject-level parameters were entered into second-level Parametric Empirical Bayes (PEB) models(46). A drug-effect model (intercept + treatment, coded psilocybin +1 / placebo −1) identified connections modulated by psilocybin, and condition-specific behavioral-association models tested whether connectivity related to the empathy outcomes showing significant psilocybin effects (positive-stimulus cognitive empathy, emotional arousal, and empathic concern; entered simultaneously and demeaned). All models underwent Bayesian model reduction and averaging, with connections interpreted at Pp ≥ 0.99 and prioritized using a tiered convergence framework (baseline connectivity, psilocybin modulation, behavioral relevance, robustness to permutation and additional bootstrap resampling; see Supplementary Methods).

### Statistical analysis

Statistical analysis was conducted in IBM SPSS Statistics 25.

Outcome variables of the MET, oxytocin concentrations, and satisfaction with life scale were analyzed using a full factorial linear mixed model (LMM) with repeated measures. Fixed effects included Treatment (psilocybin or placebo), Session (baseline, acute, follow-up), and their interaction (Treatment × Session), with Session specified as the repeated effect and a random intercept for subjects. A first-order autoregressive covariance structure (AR1) was used, which includes all available observations; participants with missing data at individual data points (MET, N = 2; oxytocin, N=22; SWLS, N=4) were therefore retained. As previous publications point to valence-specific effects of psilocybin on empathy(22, 23, 47), this analysis was repeated for negative and positive stimuli separately. Significant Treatment × Session interactions were decomposed using simple-effects analyses derived from the same model. To localize the source of each interaction, the effect of Session was tested within each treatment group (baseline vs. acute vs. follow-up), and the effect of Treatment was tested at each session; estimated marginal mean contrasts were Bonferroni-corrected for multiple comparisons within each family. Subscales on the PEQ were compared between the treatment groups using independent samples t tests; missing data (N=1) was handled by listwise deletion. A spearman correlation was used to assess the relationship between changes in oxytocin and changes in empathy.

To examine whether acute changes in empathy statistically mediated the relationship between treatment and persisting positive psychological effects, simple mediation analyses were conducted using PROCESS for SPSS (Model 4; Hayes). Treatment condition was entered as the predictor, emotional-empathy change scores (acute – baseline) measures as the mediator, and 7-day persisting psychological outcomes as the dependent variables. Mediation models were conducted separately for emotional arousal and empathic concern, and separately for each persisting outcome. These analyses were restricted to empathy variables and persisting outcomes that showed evidence of treatment-related effects in the primary analyses. Indirect effects were estimated using 5000 bootstrap samples.

For all frequentist analyses, the alpha criterion was set at a *p* value of less than .05.

**Figure 1.**
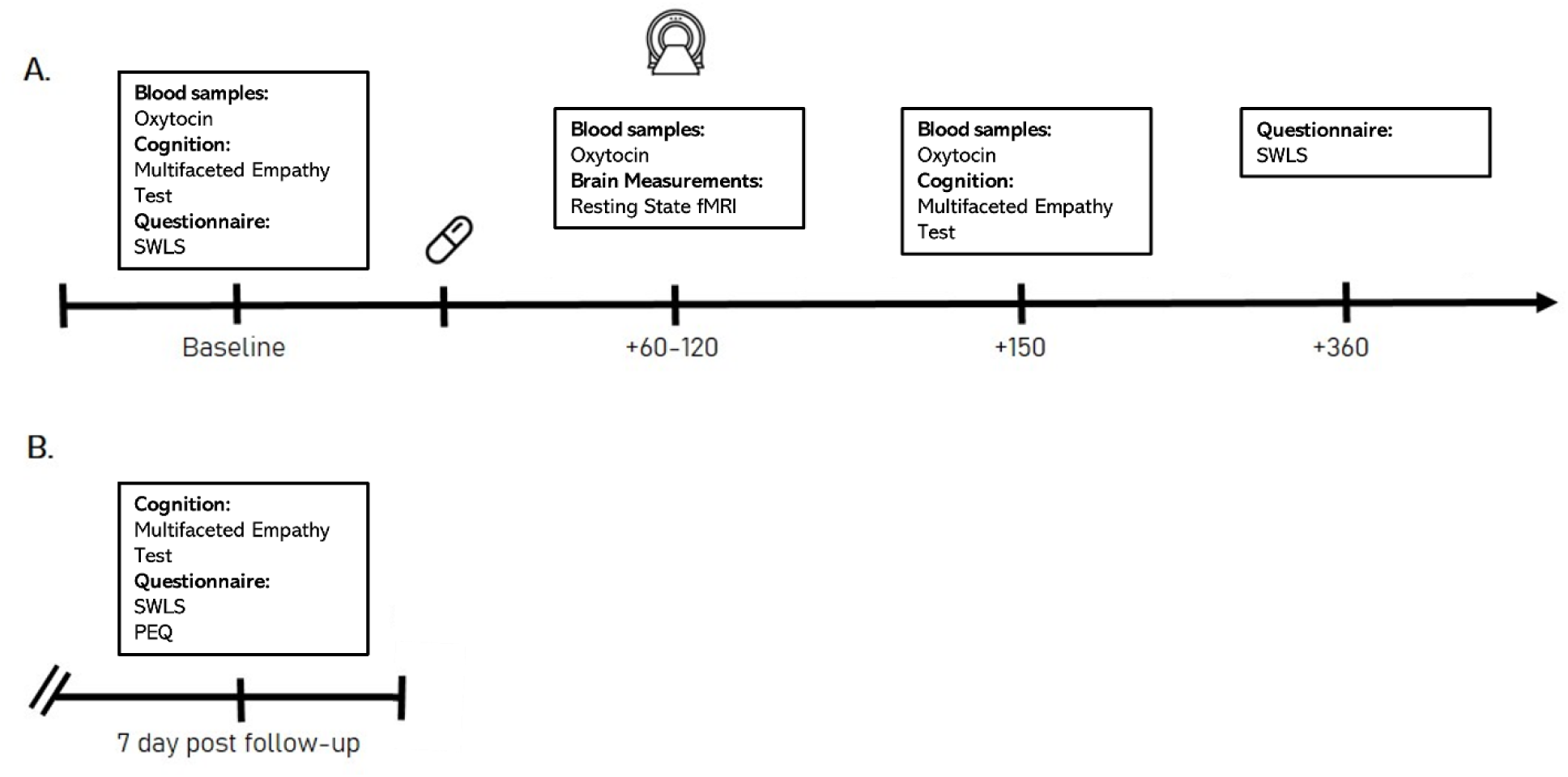
Experimental timeline. A) testing day 1, including psilocybin or placebo treatment. Timing is in minutes, relative to the treatment (psilocybin or placebo). B) testing day 2, which took place 7 days after testing day 1 (SWLS= Satisfaction with Life Scale; PEQ= Persisting Effects Questionnaire)

## Results

Demographic information has been previously published elsewhere(31–33). The psilocybin group (n = 30) and the placebo group (n = 30) did not differ in respect to demographic variables, such as sex, age, and drug use history (Table S1).

### Psilocybin altered cognitive and emotional components of empathy acutely, but not 7 days later

#### Cognitive Empathy

Cognitive empathy showed a significant Session × Treatment interaction (p = .040; Table S2). Decomposition showed that the psilocybin group decreased from baseline to the acute session (Δ = −1.71, 95% CI [−3.08, −0.34], p = .009), whereas the placebo group did not change (p = .455), indicating that psilocybin acutely reduced the ability to recognize emotional faces. This reduction was specific to positive emotions (positive-valence interaction p = .007; negative-valence p = .942; Table S2), where the psilocybin group decreased from baseline to acute (Δ = −1.45, 95% CI [−2.57, −0.33], p = .007) with no change in placebo (p = .470) (Figure 2).

**Figure 2.**
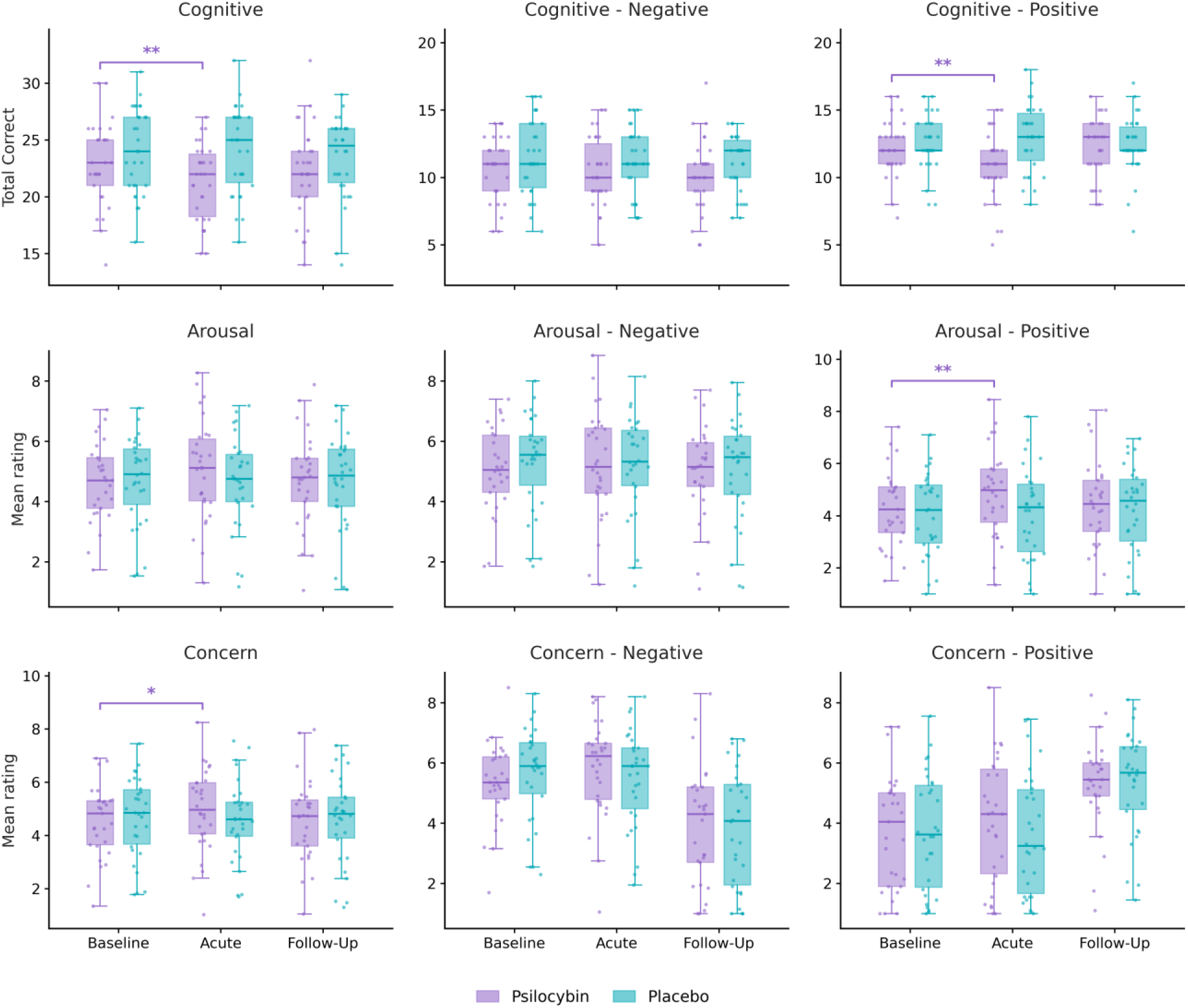
Effects of psilocybin on cognitive and emotional components of empathy across sessions. Boxplots and individual data points are shown for cognitive empathy, emotional arousal, and empathic concern, as well as for their negative and positive stimulus subsets, at baseline, acute, and follow-up sessions in the psilocybin and placebo groups. Boxes represent the interquartile range, center lines indicate the median, whiskers extend to 1.5× the interquartile range, and points represent individual participants. Means for each empathy outcome for each group and session are found in Table S3. Brackets indicate a significant within-group change from baseline to the acute session in the psilocybin group, derived from simple-effects decomposition of the Session × Treatment interaction (Bonferroni-adjusted); the placebo group showed no significant change across sessions. * indicates p < 0.05, ** indicates p < 0.01.

#### Emotional arousal

There was no interaction on overall emotional arousal (p = .175), but a significant Session × Treatment interaction on positive emotions (p = .023; negative-valence p = .587; Table S2). Here the psilocybin group increased in arousal from baseline to the acute session (Δ = 0.59, 95% CI [0.15, 1.03], p = .005), whereas placebo was unchanged (p = .696) (Figure 2).

#### Empathic concern

Empathic concern showed a significant Session × Treatment interaction (p = .038; Table S2), driven by the psilocybin group, which increased from baseline to the acute session (Δ = 0.47, 95% CI [0.08, 0.85], p = .012) and returned toward baseline at follow-up, whereas placebo did not change (p = .819). No interactions were observed for the valence-specific subscales (positive p = .294; negative p = .273) (Figure 2).

### Acute psilocybin-induced changes in empathic concern, but not emotional empathy, mediated positive persisting psychological changes

We next examined whether acute empathy changes mediated the relationship between treatment and persisting positive psychological effects. First, it was determined whether psilocybin induced positive persisting psychological effects. On the Satisfaction with Life Scale, there was no significant Session*Treatment interaction (F_2, 69.91_= 1.96, *p* = 0.14). Mean (SD) scores for the psilocybin group were 26.70 (0.99) at baseline, 27.70 (1.05) at the acute session, and 27.80 (1.05) at follow-up, compared with 27.10 (0.97), 26.76 (1.09) and 27.43 (0.99) respectively, in the placebo group.

On the Persisting Effects Questionnaire, it was found that psilocybin increased self-ratings of positive changes in attitudes about life (Mean (SD) psilocybin: 19.60 (14.87), placebo: 5.55 (8.90), *t*(57) = 4.38, *p* < 0.001), positive changes regarding attitudes about self (psilocybin: 19.60 (14.87), placebo: 4.00 (8.03), *t*(57) 4.12, *p* < 0.001), positive mood changes (psilocybin: 10.30 (10.42), placebo: 2.27 (4.21), *t*(57) = 3.90, *p* < 0.001) and positive changes in relationships (psilocybin: 10.53 (10.08), placebo: 2.38 (2.93), *t*(57) = 3.80, *p* < 0.001).

Next, mediation analyses were conducted to examine whether acute changes from baseline in emotional arousal for positive stimuli or empathic concern mediated the relationship between treatment condition and persisting positive psychological effects. In these mediation models, greater acute increases in both emotional arousal and empathic concern were positively associated with all four persisting outcomes (Table 1). However, the bootstrapped indirect effects were significant only for empathic concern — across life, self, mood, and social domains — whereas those for emotional arousal all included zero. Thus, acute empathic concern, but not emotional arousal, showed evidence of mediating the relationship between psilocybin and persisting positive psychological change.

**Table 1.**
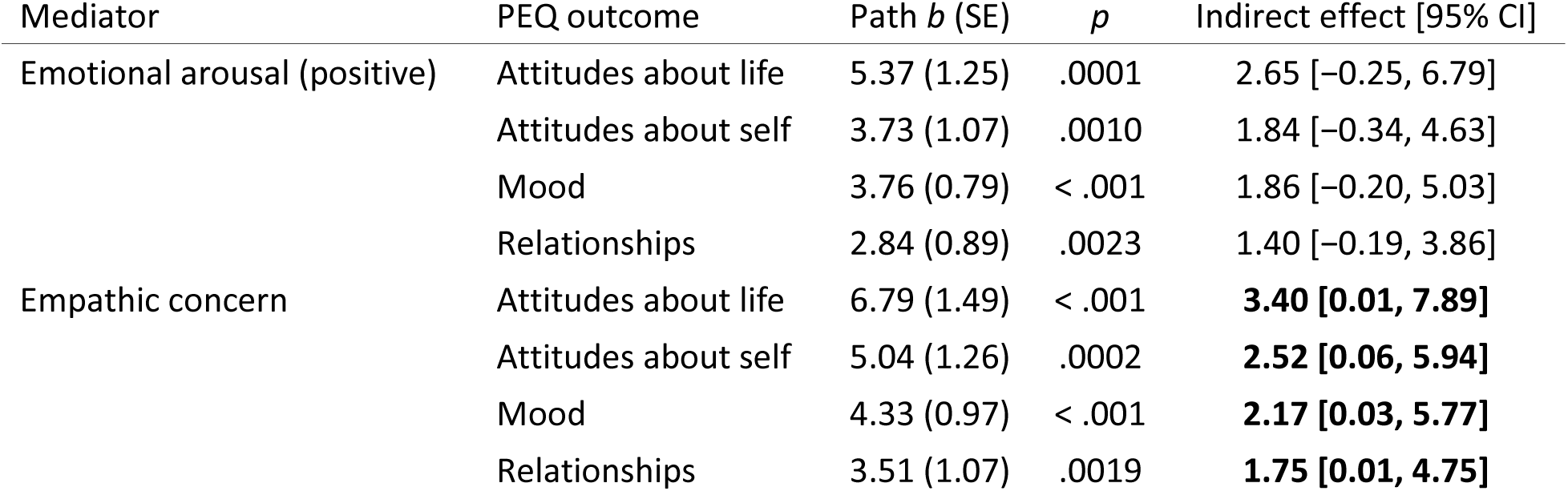
Mediation of persisting positive psychological effects by acute changes in empathy. Path *b* is the association between the acute change in each mediator (baseline-to-acute) and each Persisting Effects Questionnaire (PEQ) outcome. Indirect effects are bias-corrected bootstrap estimates (5,000 resamples) of the treatment → mediator → outcome path. Bold indirect effects have 95% confidence intervals excluding zero.

### Psilocybin increased concentrations of oxytocin, but these did not correlate with changes in empathy

Analysis revealed a significant Session*Treatment interaction effect on oxytocin concentrations (F_2,75.82_= 6.01, *p* < 0.00). Compared to placebo, psilocybin acutely increased concentration levels 80 minutes post-administration. Mean (SD) scores for the psilocybin group were 8.98 (3.90) at baseline, 10.85 (4.45) at 80 minutes post administration, and 6.77 (3.05) at 150 minutes post administration, compared with 5.12 (1.60), 5.05 (1.36) and 7.06 (1.84), respectively, in the placebo group.

There were no significant Spearmans correlations between changes in any of the MET outcome variables which showed a treatment effect, and changes in oxytocin concentrations (all *p* > 0.33).

### Psilocybin-induced changes in effective connectivity mapped onto dissociable effects on cognitive and affective empathy

We next tested whether psilocybin-induced changes in empathic responding were associated with altered directed interactions within an empathy-relevant network. Psilocybin altered effective connectivity within this network in a manner that was behaviorally relevant to empathic responding (Fig 3). Using the tiered convergence framework (see Supplementary Methods), three between-region connections met the criteria for 1) baseline connectivity, 2) psilocybin modulation and 3) behavioral relevance rSTG → lSTG, lParaHip → lSTG, and lParaHip → rParaHip. Across these connections, psilocybin was associated with weakened effective connectivity, reflecting reduced excitatory influence or increased inhibitory influence, depending on the sign of the baseline connection.

**Figure 3.**
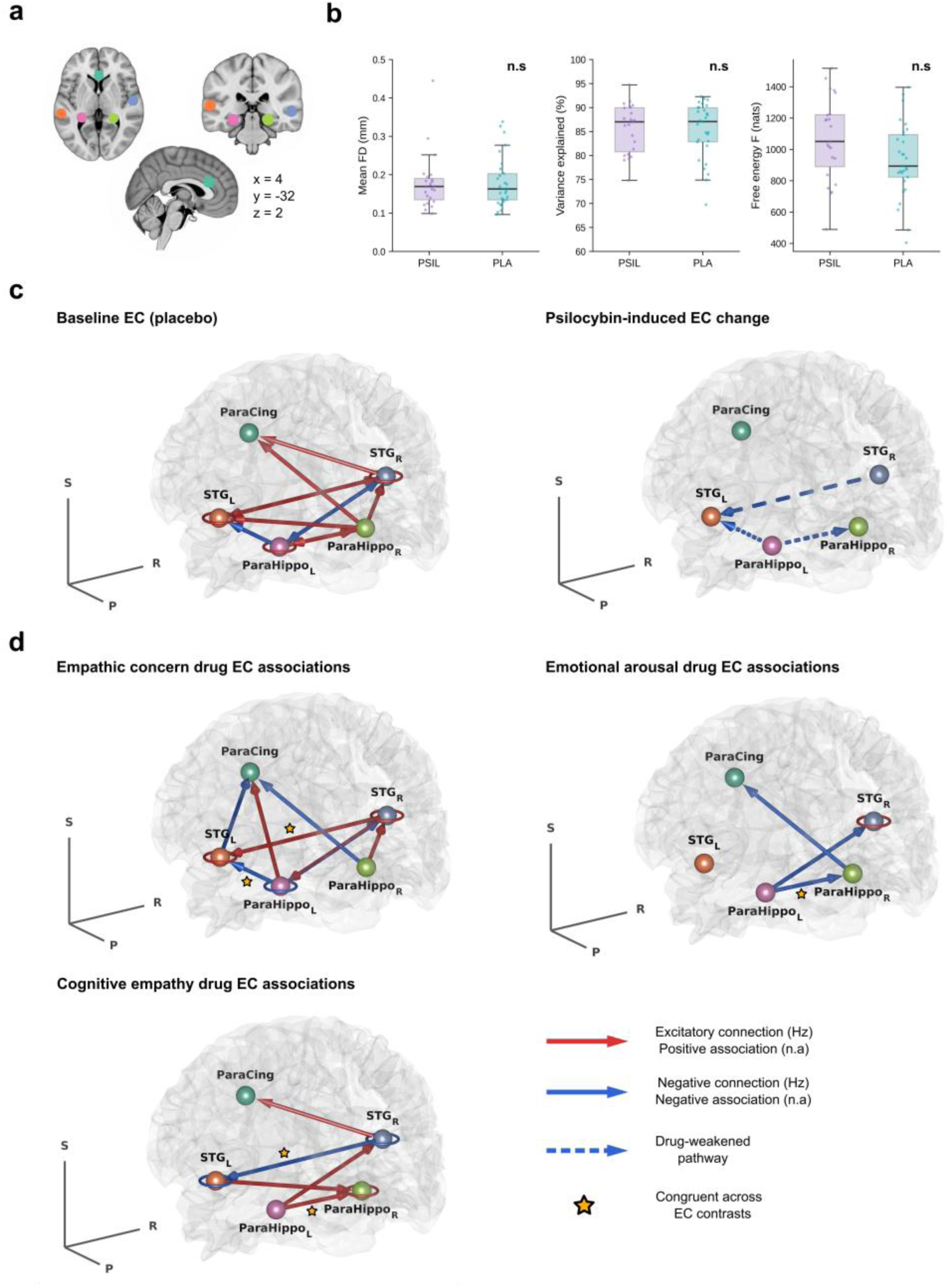
Dynamic causal modelling of effective connectivity (EC) with an empathy network and its association with empathic responding under psilocybin. A) Five regions of interest, paracingulate cortex (ParaCing), left and right superior temporal gyrus (lSTG, rSTG), and left and right parahippocampal cortex (lParaHip, rParaHip), were defined from Neurosynth meta-analytic association maps as 8-mm spherical seeds centered on each peak coordinate (see Methods for exact MNI coordinates). Acute rsfMRI data acquired approximately 102 min post administration were fitted using spectral DCM, which estimates directed EC between regions by modelling local haemodynamics and neural population dynamics. B) Derived DCM quality control measures. QC metrics: mean FD = average framewise displacement in mm during fMRI acquisition, variance explained = % variance in regional responses captured by the DCM fit (higher is better); Free energy = per-subject variational free energy following DCM model inversion (higher reflects greater model evidence). Two-sample t-tests did not identify significant differences across all measures (p > 0.05). C) Effective connective maps from a hierarchical PEB model are shown. Baseline EC (placebo): posterior expectation of effective connectivity at the intercept of the drug-effect PEB, in Hz; red indicates positive (excitatory) and blue negative (inhibitory) influences. Psilocybin-induced change: posterior expectations of the [psilocybin − placebo] contrast, in Hz; red indicates an increase (increased excitation or decreased inhibition) and blue a decrease (decreased excitation or increased inhibition) relative to placebo, regardless of the baseline sign. Self-connections at baseline are inherently inhibitory D) MET behavioral associations of drug-induced EC changes. Regression coefficients from the combined MET PEB linking each connection’s value to each MET empathy domain under psilocybin; red = positive Ep (more excitatory connection or stronger self-inhibition co-varies with higher empathic concern) and blue = negative Ep. Between-region connections are rendered as cylinders with a cone arrowhead at the target; Dashed cylinders denote attenuated connections. In spectral DCM, the self-connections are log-scaling factors on an intrinsically inhibitory term. For self-connections (rendered as rings), baseline positive Ep (red) therefore reflect stronger self-inhibition. Connections are displayed only when the posterior probability that the corresponding parameter is non-zero exceeds 0.99, applied separately for each map; therefore, a connection can be absent in one map yet present in another if there is stronger evidence in the latter. ★ marks behavioural associations consistent with EC change contrasts. All maps satisfy a Bayesian-local-FDR multiplicity guarantee (mean q < 0.001 per matrix). The reported effects additionally survived a non-parametric permutation null on the BMA effect magnitude (pperm < 0.05) and a subject-bootstrap stability criterion (Pp ≥ 0.99 retained in ≥ 50 % of resamples). The full DCM matrices (including placebo associations) and corresponding values are provided in the Supplementary Materials.

Under psilocybin, cross-hemispheric STG coupling from rSTG to lSTG was positively associated with empathic concern such that less excitatory effective connectivity was related to lower empathic concern, further surviving bootstrapping-based validation (p = 0.038, see Supplementary). This same connection showed an inverse association with cognitive empathy, such that less excitatory effective connectivity was related to greater cognitive empathy. lParaHip → lSTG connectivity was negatively associated with empathic concern, indicating that more inhibitory influence from left parahippocampal cortex to lSTG was associated with greater empathic concern. The lParaHip → rParaHip connection was associated with both affective and cognitive components: greater inhibitory effective connectivity was related to higher emotional arousal and to higher cognitive empathy for positive stimuli. Additional connections otherwise not modulated at the group level by psilocybin showed robust state-dependent associations with empathic responding across the validation criterion (see Supplementary, Figures S2/S3). Together, these findings suggest that psilocybin reorganized effective connectivity within an empathy-relevant network such that cognitive and affective empathy were associated with opposing patterns of directed interactions.

## Discussion

The present findings indicate that psilocybin does not produce a global enhancement of empathic ability, but instead induces a selective rebalancing of empathic processing, with dissociable effects across cognitive and affective components. Acutely, psilocybin reduced cognitive empathy while increasing empathic concern and emotional arousal for positive stimuli; none of these effects persisted at seven days. However, acute increases in empathic concern — but not emotional arousal — showed significant indirect effects on multiple domains of positive psychological change at follow-up, suggesting that the acute rebalancing may carry downstream relevance even without direct persistence. Psilocybin also acutely increased circulating oxytocin, but these changes were unrelated to empathy. Instead, empathic responding was associated with altered effective connectivity within an STG-centered network, suggesting that superior temporal dynamics may represent one neural substrate through which psilocybin differentially affects cognitive and affective empathy.

### Acute effects of psilocybin on cognitive and affective components of empathy

Psilocybin reduced cognitive empathy selectively for positive stimuli, leaving recognition of negative states unchanged. This valence-specificity argues against a generalized performance impairment and instead indicates that psilocybin reduced the precision of explicit emotion identification in a stimulus-dependent manner. The finding adds to a mixed literature in which psychedelics have been reported to impair recognition of negative stimuli, enhance recognition of positive stimuli, or leave cognitive empathy unchanged(13, 20, 21, 29, 47), suggesting that cognitive empathy may be more vulnerable to compound, dose, and timing than affective components, which more consistently increase.

In contrast, psilocybin increased empathic concern across valences while increasing emotional arousal only for positive stimuli. This pattern indicates a shift away from precise explicit emotion identification and toward affective engagement with others’ emotional states, and shows that concern and arousal were not affected identically: psilocybin may broadly enhance other-oriented concern while more selectively amplifying affective responsiveness to positive social-emotional cues.

The clinical implications of this dissociation are likely to be context-dependent. In conditions characterized by reduced emotional responsiveness, social withdrawal, or rigid self-referential processing — as seen in major depression — an acute shift toward greater affective engagement and other-oriented concern may be functionally adaptive, at least transiently (48). In conditions involving affective overreactivity or impaired mentalizing, such as borderline personality disorder or some presentations on the autism spectrum, a simultaneous increase in affective engagement and decrease in social-cognitive accuracy could instead exacerbate existing difficulties (49, 50). The present findings should therefore not be interpreted as evidence that psilocybin improves social functioning, but as evidence that it acutely alters the balance between dissociable components of empathic responding — with consequences that depend on clinical presentation, interpersonal context, and the circumstances in which the drug state unfolds. This context-sensitivity is directly relevant to psychedelic-assisted psychotherapy: heightened relational sensitivity may facilitate emotional disclosure and openness, while reduced cognitive precision increases the potential for misattribution, projection, or misattunement(51).

### Persisting effects of psilocybin on cognitive and affective components of empathy

All empathy-related effects were limited to the acute state, with no changes at seven days. This contrasts with the few prior studies reporting persisting increases, which were conducted in explicitly social settings such as ceremonies or retreats(22, 23), or within a therapeutic framework(24). Notably, our dose (0.17 mg/kg) was comparable to the therapeutic study (0.215 mg/kg), suggesting dose alone does not explain the absence of persisting effects. Persisting empathy-related changes may therefore not unfold through drug effects within an isolated individual, but may depend on the relational context in which acute social-affective changes are experienced and reinforced. Because the present study involved no extended social contact, the acute effects appear pharmacologically driven while their persistence may be more dependent on interpersonal context.

### Acute changes in affective empathic responding and persisting positive psychological effects

Acute increases in empathic concern showed significant indirect effects across multiple domains of persisting positive change — attitudes about life and self, mood, and relationships — suggesting a possible mediating role in the relationship between psilocybin and later positive outcomes, consistent with naturalistic studies linking post-acute empathy increases to greater wellbeing(22, 23). Empathic concern may be especially relevant because it reflects an other-oriented stance of care rather than affective activation alone; concern and compassion are often used interchangeably(52), and concern for others is positively associated with wellbeing (53). Such an other-oriented stance may be more likely to interrupt rigid self-focus and promote prosociality, connectedness, and more positive appraisals of self and relationships, all proposed as contributors to the enduring effects of psychedelics(54–56). Empathic concern may thus contribute to later change indirectly, by enabling broader self-related and interpersonal processes that unfold after the acute state(24).

### Biological correlates of psilocybin-induced changes in empathy

Psilocybin acutely increased circulating oxytocin concentrations, but these changes were not associated with changes in cognitive empathy, emotional arousal, or empathic concern, indicating that the acute empathy effects cannot be explained by peripheral oxytocin alone. This is consistent with evidence that LSD-induced increases in emotional empathy persist even when ketanserin blocks oxytocin increases, suggesting effects that are independent of peripheral oxytocin and perhaps of 5-HT2AR engagement(29).

At the neural level, empathic responding under psilocybin was related to directed interactions within an STG-centered network. Superior temporal regions are central to social perception and mental-state inference(41, 57), while parahippocampal regions may contribute contextual and associative information that gives social-emotional cues meaning by linking perception to context, memory, or self-referential experience(43, 58). Cross-hemispheric coupling from right to left STG was associated with empathic concern and cognitive empathy in opposite directions, suggesting that interhemispheric superior temporal dynamics may reflect a trade-off between affective engagement and explicit emotion identification rather than supporting empathic responding uniformly. The parahippocampal connections, by contrast, may reflect changes in how socially relevant cues are integrated with contextual information during empathic responding. This fits models in which empathy depends on both bottom-up social-perceptual and top-down integrative processes(59, 60): psilocybin increased affective engagement while reducing accuracy in explicitly identifying positive states, and because positive empathic processing may recruit additional regulatory mechanisms(60, 61), psilocybin may have increased the affective salience of positive social cues while reducing the precision of the integrative processes needed to identify the specific emotion depicted.

### Limitations and conclusions

Several limitations should be noted. The neuroimaging was resting-state while empathy was assessed outside the scanner, so the DCM findings should be interpreted as neural correlates of empathic responding rather than task-evoked or causal mechanisms. A key next step will therefore be live, interactive assessment, such as dual-brain hyperscanning during real-time social exchange, to capture the neural dynamics of empathic responding as they occur under a psychedelic(62). The persisting outcomes relied on self-report PEQ ratings, which may be vulnerable to retrospective or expectancy bias. And although the mediation analyses suggest acute empathic concern may contribute to later positive change, the indirect effects were modest and statistical mediation cannot establish causality.

Despite these limitations, the findings show a coherent pattern in which psilocybin acutely shifted empathic responding away from explicit emotion identification and toward affective, other-oriented responding. These effects did not persist at seven days, but acute increases in empathic concern were associated with later positive psychological change, and effective-connectivity findings suggest psilocybin reorganizes the acute balance between empathic concern, emotional arousal, and cognitive empathy within an STG-centered network. Together, these results indicate that psilocybin-induced changes in empathy are not explained by peripheral oxytocin alone, and underscore the importance of acute relational and social-affective processes in psychedelic research and therapy.

## Acknowledgements

This project received funding from Amanda Feilding and the Beckley Foundation. NLM is financially supported by the Dutch Research Council (NWO, grant number VI.Veni.231G.011). Thank you to all participants for the time, effort and trust, to Cees van Leeuwen for medical supervision, and to all interns for their help in participant recruitment, screening, and data collection. During the preparation of this work, the author(s) used Claude (Anthropic) to refine, edit, and format human-written text; and to produce code used to generate figures. All AI-assisted output was reviewed, verified, and edited by the author(s), who take full responsibility for the content of the publication.

## Author contributions

Conceptualization: NLM, KPCK, JGR. Data acquisition: NLM, JTR. Analysis: NLM, PM, RDLTF. Original draft: NLM, PM. Writing-Review & Editing: all authors. Funding Acquisition: NLM & JGR.

## Data availability

Data can be made available upon reasonable request to N.L.M. and J.G.R. and a data use agreement executed with Maastricht University.

## Supplementary Materials

### Methods

#### Participants

Participants were recruited through advertisements around Maastricht University and internet forums in the Netherlands. Inclusion criteria were: age, 18-40 years; previous experience with a psychedelic drug, but not within the past 3 months; normal weight, body mass index between 18 and 28 kg/m2; free from psychotropic medication; good physical health, including absence of major medical, endocrine, and neurological conditions; and written informed consent. Exclusion criteria were: history of drug abuse or addiction; pregnancy or lactation; health issues including hypertension (diastolic >90 and systolic >140), cardiac dysfunction, and liver dysfunction; current or history of psychiatric disorders; previous experience of serious side effects to psychedelics; and MRI contraindications. Before inclusion, subjects answered medical questionnaires about their health and drug use, and were screened and examined by a study physician, who checked for general health, conducted a resting ECG, and took blood and urine samples in which hematology, clinical chemistry, urine, and virology analyses were conducted. Participant demographic data can be found in the previous publication(31).

Psilocybin (powder) was obtained from GH Pharm GmbH, Frankfurt, Germany. A permit for obtaining, storing, and administering psilocybin was obtained from the Dutch Drug Enforcement Administration. Participants were financially compensated for their participation in the study.

#### Randomization and blinding

An experimenter who was not responsible for treatment randomization or preparation, recruited all participants. A separate experimenter, who did not come in direct contact with the subjects, allocated treatment in a completely random order, using a random number generator. This latter experimenter was also responsible for preparing the treatment (psilocybin or placebo), and giving the treatment to the data collector in a closed cup. The closed cup ensured that neither the data collector or the participant would be unblinded as to what treatment the participant was receiving. Only following completion of all data collection, was the study fully unblinded.

#### Sample size calculation

The overall study sample size was determined in regards to the imaging data. Power analysis for imaging data is complex. Because data analysis of imaging data consists of a very large number of non-independent multiple comparisons, a proper power analysis with correction for multiple comparisons requires computer simulation. In addition, statistical power is not the same (i.e., homogenous) over the brain, but varies across brain regions. The most common approach in the is to estimate the number of subjects from previous empirical experience in the field. Previous MRS and FC studies have employed 10-20 subjects for showing significant drug effects. Based on these experiences we expected that we would have no problem to demonstrate drug effects on MRS and FC in 30 participants in each drug condition.

#### Procedure

Participants were familiarized with the test day procedures on a separate training day prior to the treatment conditions. Participants were instructed to refrain from drug use, including psychedelic drugs (≥ 3 months), MDMA/ecstasy (≥ 14 days), alcohol (≥24 hours), and all other drugs of abuse (≥7 days) prior to their first testing day, and to remain sober until completion of the follow-up testing day (7 days later). Additionally, participants were asked to refrain from caffeine and nicotine use the day of the test day.

On arrival of a test day, absence of drug and alcohol use was assessed via a urine drug screen and a breath alcohol screen. An additional pregnancy test was given if participants were female. If all tests were found to be negative, participants were allowed to proceed, and a venal catheter was placed, in order to take blood samples throughout the testing day. Before administration of treatment, a baseline blood sample was taken and baseline vital signs (blood pressure and heart rate) were measured. After measurements, the treatment was administered orally, in a closed cup containing bitter lemon (placebo) or bitter lemon and psilocybin (powder; 0.17 mg/kg psilocybin). Bitter lemon was used in order to conceal any potential taste of psilocybin. After 40 minutes, participants were placed in the MRI scanner, where resting state scans and magnetic resonance spectroscopy were performed throughout a 1 hour time window. When they were taken out of the MRI scanner, they returned to the laboratory room where they completed the creativity tests. At the end of the test day (approximately 6 hours after treatment administration), participants were asked to complete measures of retrospective subjective high. Participants stayed under supervision until the testing day was complete, and the researcher deemed they were fit to go home. Participants then returned 7 days later, and were screened again for drug and alcohol use, and if applicable pregnancy, via the aforementioned methods. If the tests were found to be negative, they were allowed to proceed. If they were positive, participants were sent home and excluded from the study (n=0).

#### Satisfaction with life scale

Individuals answer each item on a Likert scale ranging from 1 (strongly disagree) to 7 (strongly agree). The total score is then obtained by summing the ratings form each item. Sample items include “In most ways my life is close to my ideal” and “If I could live my life over, I would change almost nothing.” The minimum possible score is 5, and the maximum possible score 35, with a score of 5–19 defined as dissatisfied to below average life satisfaction, 20–24 defined as average life satisfaction, and 29–35 defined as high to very high life satisfaction (36). The scale has previously been shown to be a valid and reliable measure of life satisfaction (36).

#### Persisting effects questionnaire

The questionnaire included four additional questions (see <u>Griffiths et al.</u> <u>2006</u> for details): 1) *How personally meaningful was the experience?* (rated from 1 to 8, with 1= *no more than routine*, and 8= *the single most meaningful experience of my life*); 2) *Indicate the degree to which the experience was spiritually significant to you?* (rated from 1 to 8, using the same encoding as the first question); 3) *How psychologically challenging were the most psychologically challenging portions of the experiences?* (rated from 1 to 8, using the same encoding as the first question); 4) *How personally psychologically insightful to you were the experiences?* (rated from 1 to 8, using the same encoding as the first question).Full results from this questionnaire can be found in the previous publication(33)

#### Oxytocin

A fluorescence immunoassay kit (Phoenix Pharm. Inc., Burlingame, CA) was used to measure serum oxytocin concentrations in accordance with the manufacturer’s instructions. In non-heparinized tubes, a 2-mL sample was taken for oxytocin assay. Samples were centrifuged at 4 °C for 10 minutes at 3500 rpm. Before analysis, the serum was extracted and refrigerated at −80 °C. In short, a fluorimetric EIA analysis was conducted in accordance with protocol after a first extraction step utilizing C18 columns.

### DCM Methods

#### Neuroimaging acquisition

Participants underwent structural MRI (50 min post-treatment) and eyes-open rsfMRI (102 min post) during peak subjective drug effects. Images were acquired on a MAGNETOM 7T MR scanner. At rest, 258 whole-brain EPI volumes were acquired (TR = 1400 ms; TE = 21 ms; field of view (FOV) = 198 mm; flip angle = 60°; oblique acquisition orientation; interleaved slice acquisition; 72 slices; slice thickness = 1.5 mm; voxel size = 1.5 × 1.5 × 1.5 mm). During scanning, participants were shown a black cross on a white background and instructed to look at the cross while attempting to clear their mind and lie as still as possible. Detailed information regarding image acquisition was previously reported (Mason et al., 2020)

#### MRI preprocessing and quality control

Preprocessing was performed in SPM12 using a previously validated effective-connectivity pipeline and included dummy-volume removal, slice-timing correction, realignment, structural co-registration, segmentation, normalization, and spatial smoothing with a 6-mm full-width-at-half-maximum Gaussian kernel(63). Regional time series were extracted from five empathy-relevant regions of interest: paracingulate cortex, left and right superior temporal gyrus, and left and right parahippocampal cortex. Nuisance regressors included six rigid-body motion parameters, cerebrospinal fluid signal, and white-matter signal. Low-frequency drifts were removed using a 128-s high-pass filter, and global signal regression was not applied. Participants with mean framewise displacement greater than 0.5 mm were excluded before DCM analysis. After exclusion, mean framewise displacement did not differ between the psilocybin and placebo groups.

#### Spectral dynamic causal modeling

For each participant, a fully connected spectral DCM was specified using the five selected regions of interest. The model included all endogenous connections among regions and no exogenous inputs. Spectral DCM estimates directed effective connectivity by fitting a biophysical state-space model to the cross-spectral density of the BOLD signal. This yields estimates of directed coupling rates between regions, as well as estimates of each region’s intrinsic self-inhibition. Off-diagonal connections were modeled on their native scale. Self-connections are log-scaled and are inherently inhibitory, such that positive parameter estimates reflect increased self-inhibition, and negative estimates reflect disinhibition. Participants with DCM explained variance below 60% were also excluded, yielding a final DCM sample of 52 participants.

#### Parametric empirical Bayes models

Subject-level DCM parameters were entered into second-level Parametric Empirical Bayes models. Two classes of PEB models were estimated. First, a drug-effect PEB model tested group-level effects of psilocybin on effective connectivity. The design matrix included an intercept and treatment condition, coded as psilocybin = +1 and placebo = −1. This model was used to identify baseline effective connectivity at the group level and connections modulated by psilocybin. Second, condition-specific behavioral-association PEB models tested whether individual differences in effective connectivity were associated with empathic responding. Separate models were estimated for psilocybin and placebo groups. Behavioral variables were restricted to empathy outcomes that showed significant psilocybin effects in the primary behavioral analyses (positive-stimulus cognitive empathy, empathic concern, and empathic arousal). These variables were entered simultaneously and demeaned, so that each association reflected the unique contribution of that empathy component while accounting for the others. All PEB models underwent Bayesian model reduction, and Bayesian model averaging over the 256 best-fitting reduced models (default). Effects were interpreted using a posterior probability threshold of Pp ≥ 0.99.

#### Multiplicity-aware Bayesian inference

To further account for multiplicity across connection-level inferences, complementary multiplicity-aware quantities were derived using an empirical-Bayes two-groups framework. For each connection, the Bayesian local false-discovery rate was computed as:

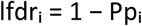

where Ppᵢ is the posterior probability that connection i is nonzero. This value corresponds to the posterior probability that the connection is null given the observed data. Bayesian q-values were then computed at the selection threshold τ as the mean local false-discovery rate across all connections exceeding the threshold:

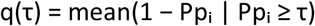

The q-value provides an estimate of the expected proportion of false positives among the selected connections.

#### Tiered convergence framework

To further distinguish findings from broader network effects, effective-connectivity results were interpreted using a tiered convergence framework. Connections were evaluated according to three criteria:

1. Baseline connectivity: the connection was present at the group level, defined by Pp ≥ 0.99 for the intercept of the drug-effect PEB.
2. Drug sensitivity: the connection was modulated by psilocybin, defined by Pp ≥ 0.99 for the treatment effect in the drug-effect PEB.
3. Behavioral relevance: the connection was associated with at least one empathy outcome in the behavioral-association PEB models, defined by Pp ≥ 0.99.

Connections meeting all three criteria were further interpreted as psilocybin-sensitive, behaviorally relevant at the group level.

#### Validation permutation-testing and bootstrapping

In addition to our primary Bayesian inference results, we assessed robustness using model-averaged effect sizes (Ep, following BMR. For the permutation null, rows of the questionnaire matrix were shuffled 1000 times while preserving within-subject covariance among MET subscales, and the full PEB+BMA pipeline was refitted each time. Empirical p-values were calculated as the proportion of null effects with Ep_null ≥ Ep_obs. For bootstrap stability, subjects were resampled with replacement 1000 times while preserving condition-specific sample sizes, and the full PEB+BMA pipeline was rerun for each resample. Stability was defined as the proportion of bootstrap samples in which Pp ≥ 0.99 was retained.

**Table S1.**
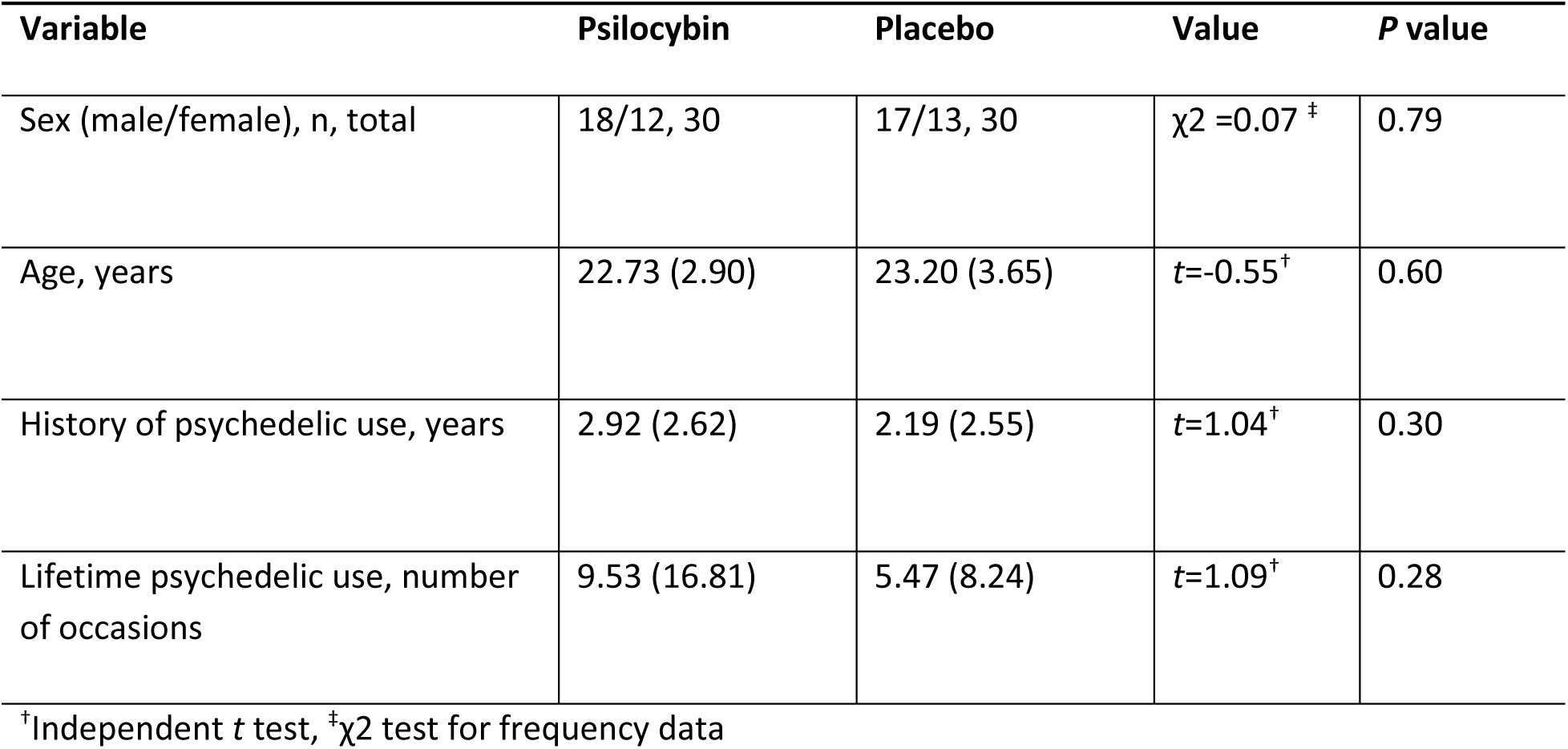
Mean subject characteristics (SD) and history of psychedelic use for healthy participants in the psilocybin and the placebo condition (N=60), as previously published. (**31, 33**).

**Table S2.**
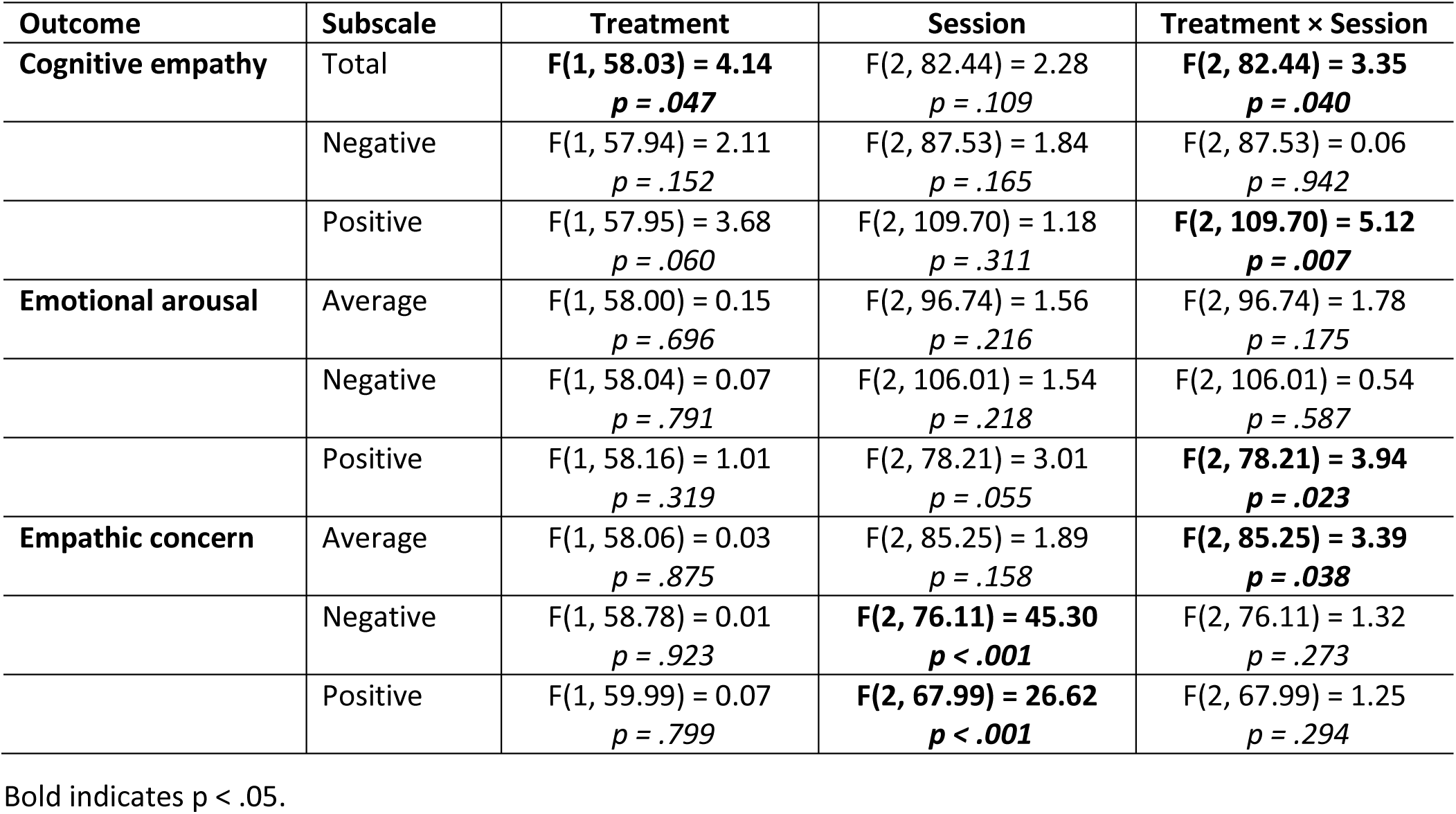
Linear mixed model results for Multifaceted Empathy Test (MET) outcomes.

| Outcome | Subscale | Treatment | Session | Treatment × Session |
| --- | --- | --- | --- | --- |
| Cognitive empathy | Total | **F(1, 58.03) = 4.14**<br>**p = .047** | F(2, 82.44) = 2.28<br>p = .109 | **F(2, 82.44) = 3.35**<br>**p = .040** |
|  | Negative | F(1, 57.94) = 2.11<br>p = .152 | F(2, 87.53) = 1.84<br>p = .165 | F(2, 87.53) = 0.06<br>p = .942 |
|  | Positive | F(1, 57.95) = 3.68<br>p = .060 | F(2, 109.70) = 1.18<br>p = .311 | **F(2, 109.70) = 5.12**<br>**p = .007** |
| Emotional arousal | Average | F(1, 58.00) = 0.15<br>p = .696 | F(2, 96.74) = 1.56<br>p = .216 | F(2, 96.74) = 1.78<br>p = .175 |
|  | Negative | F(1, 58.04) = 0.07<br>p = .791 | F(2, 106.01) = 1.54<br>p = .218 | F(2, 106.01) = 0.54<br>p = .587 |
|  | Positive | F(1, 58.16) = 1.01<br>p = .319 | F(2, 78.21) = 3.01<br>p = .055 | **F(2, 78.21) = 3.94**<br>**p = .023** |
| Empathic concern | Average | F(1, 58.06) = 0.03<br>p = .875 | F(2, 85.25) = 1.89<br>p = .158 | **F(2, 85.25) = 3.39**<br>**p = .038** |
|  | Negative | F(1, 58.78) = 0.01<br>p = .923 | **F(2, 76.11) = 45.30**<br>**p < .001** | F(2, 76.11) = 1.32<br>p = .273 |
|  | Positive | F(1, 59.99) = 0.07<br>p = .799 | **F(2, 67.99) = 26.62**<br>**p < .001** | F(2, 67.99) = 1.25<br>p = .294 |

Bold indicates p < .05.

**Table S3.** Means (SDs) of empathy scores, separated by group and session.

| **Construct** | **Baseline** | **Acute** | **Followup** | **Baseline** | **Acute** | **Followup** |
| --- | --- | --- | --- | --- | --- | --- |
| *Psilocybin* |  |  |  | *Placebo* |  |  |
| Cognitive empathy | 22.87 (3.63) | 21.22 (3.28) | 22.28 (4.00) | 24.34 (3.77) | 24.41 (3.90) | 23.64 (3.66) |
| Cognitive empathy – positive emotions | 12.27 (2.05) | 10.77 (2.50) | 12.10 (2.22) | 12.69 (2.08) | 13.12 (2.50) | 12.52 (2.20) |
| Cognitive empathy – negative emotions | 10.60 (2.30) | 10.45 (2.42) | 10.17 (2.69) | 11.66 (2.77) | 11.28 (2.47) | 11.13 (2.16) |
| Implicit emotional empathy | 4.68 (1.28) | 5.04 (1.62) | 4.69 (1.52) | 4.67 (1.43) | 4.63 (1.49) | 4.64 (1.57) |
| Implicit emotional empathy – positive emotions | 4.27 (1.43) | 4.82 (1.65) | 4.38 (1.65) | 4.04 (1.54) | 4.05 (1.71) | 4.22 (1.70) |
| Implicit emotional empathy – negative emotions | 5.10 (1.39) | 5.27 (1.77) | 5.00 (1.54) | 5.30 (1.55) | 5.21 (1.59) | 5.06 (1.68) |
| Explicit emotional empathy | 4.55 (1.32) | 4.94 (1.52) | 4.59 (1.54) | 4.67 (1.47) | 4.57 (1.48) | 4.61 (1.60) |
| Explicit emotional empathy – positive emotions | 3.75 (1.87) | 4.06 (2.03) | 5.26 (1.52) | 3.72 (1.88) | 3.64 (1.98) | 5.36 (1.69) |
| Explicit emotional empathy – negative emotions | 5.34 (1.31) | 5.82 (1.57) | 3.91 (1.94) | 5.62 (1.51) | 5.51 (1.55) | 3.86 (1.91) |

**Figure S1.**
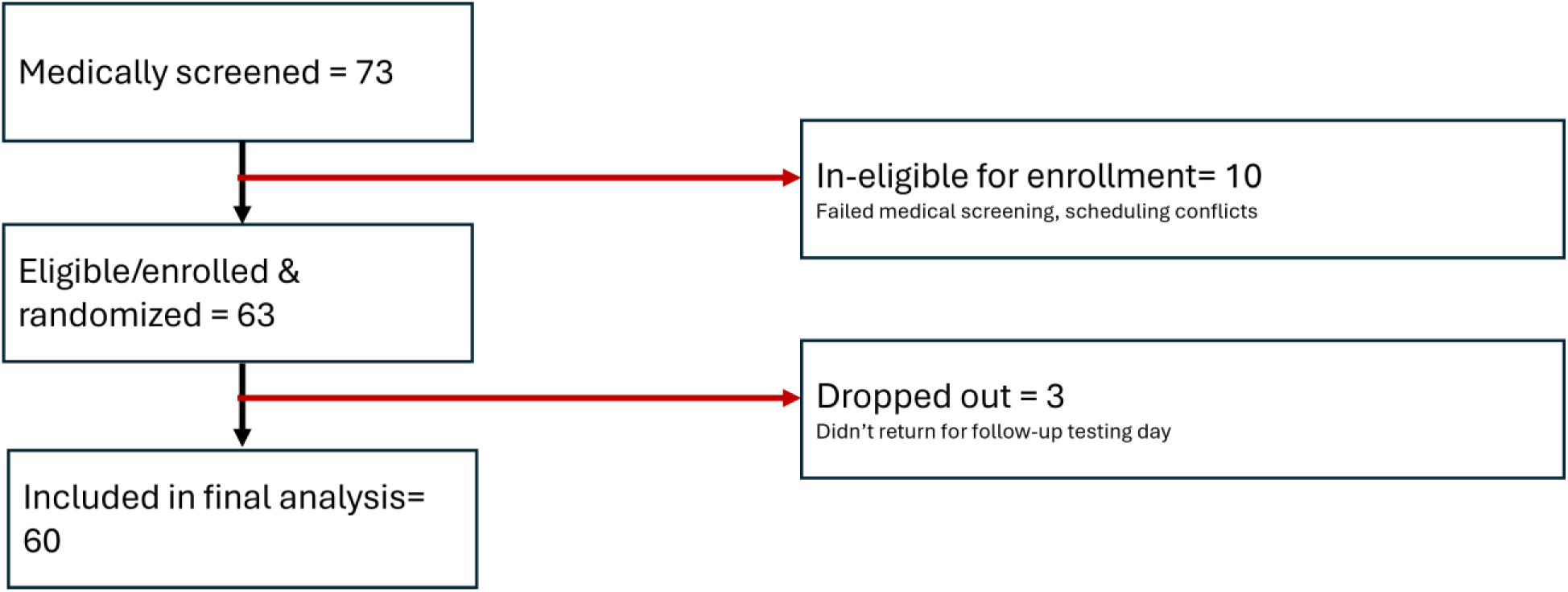
CONSORT flowchart

**Figure S2.**
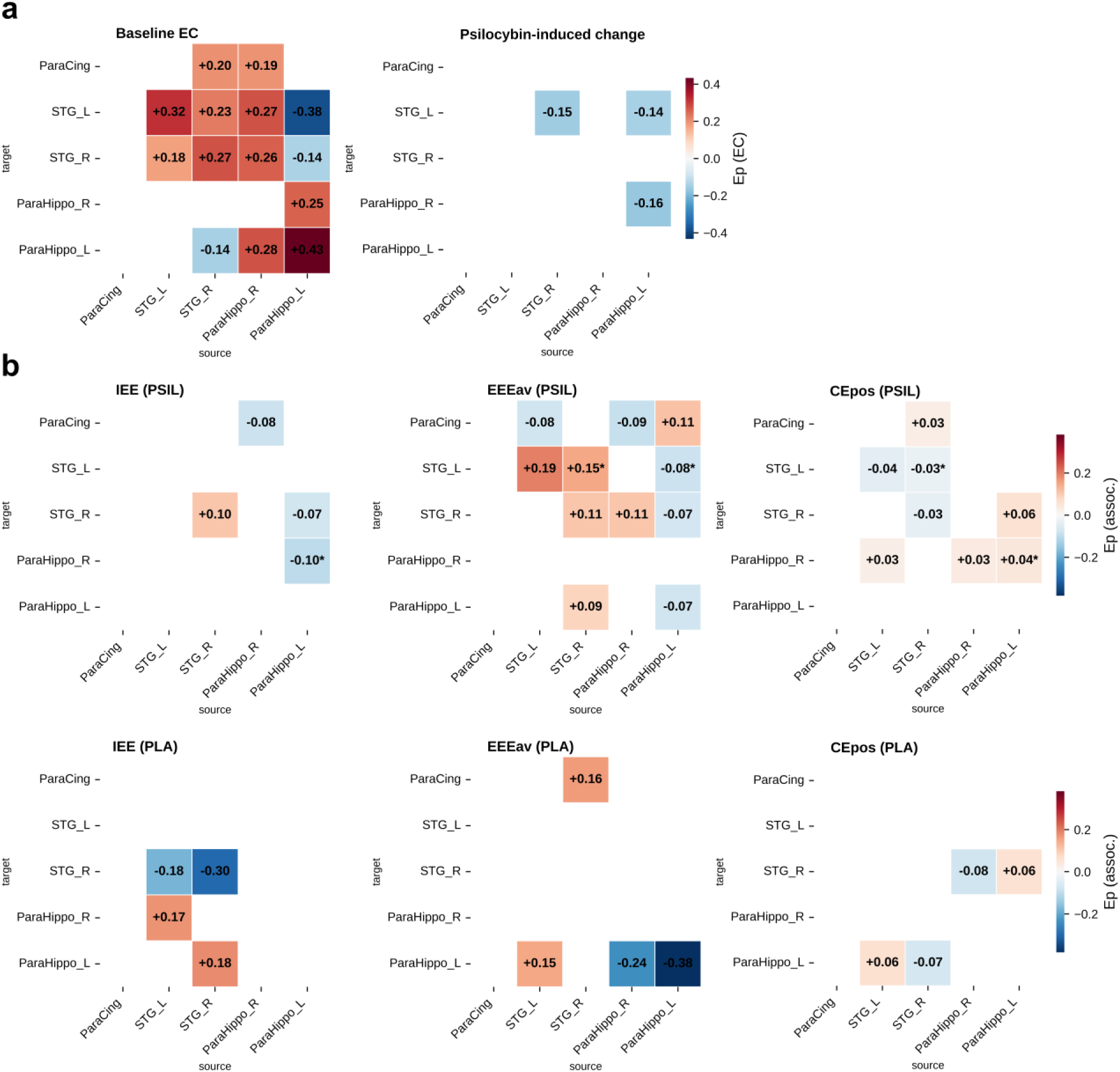
Raw DCM effective connectivity matrices. Neurosynth-derived DCM effective connective matrices. A) Effective connective matrices from a hierarchical PEB model are shown. A) baseline EC (placebo): posterior expectation of effective connectivity at the intercept of the drug-effect PEB, in Hz; red indicates positive (excitatory) and blue negative (inhibitory) influences. Psilocybin-induced change: posterior expectations of the [psilocybin − placebo] contrast, in Hz; red indicates an increase (increased excitation or decreased inhibition) and blue a decrease (decreased excitation or increased inhibition) relative to placebo, regardless of the baseline sign. C) MET behavioural associations of drug-induced EC changes. B) Regression coefficients from the combined MET PEB linking each connection’s value to each MET empathy domain under psilocybin; red = positive Ep (more excitatory connection or stronger self-inhibition co-varies with higher empathic concern) and blue = negative Ep. Connections are displayed only when the posterior probability that the corresponding parameter is non-zero exceeds 0.99, applied separately for each map; therefore a connection can be absent in one map yet present in another if there is stronger evidence in the latter. * marks behavioural associations consistent with EC change contrasts. In spectral DCM the self-connections are log-scaling factors on an intrinsically inhibitory term. Self-connections at baseline showing positive Ep (red) therefore reflect stronger self-inhibition. All maps satisfy a Bayesian-local-FDR multiplicity guarantee (mean q < 0.001 per matrix). IEE = emotional arousal, EEEav = empathic concern, CEpos = cognitive empathy

**Figure S3.**
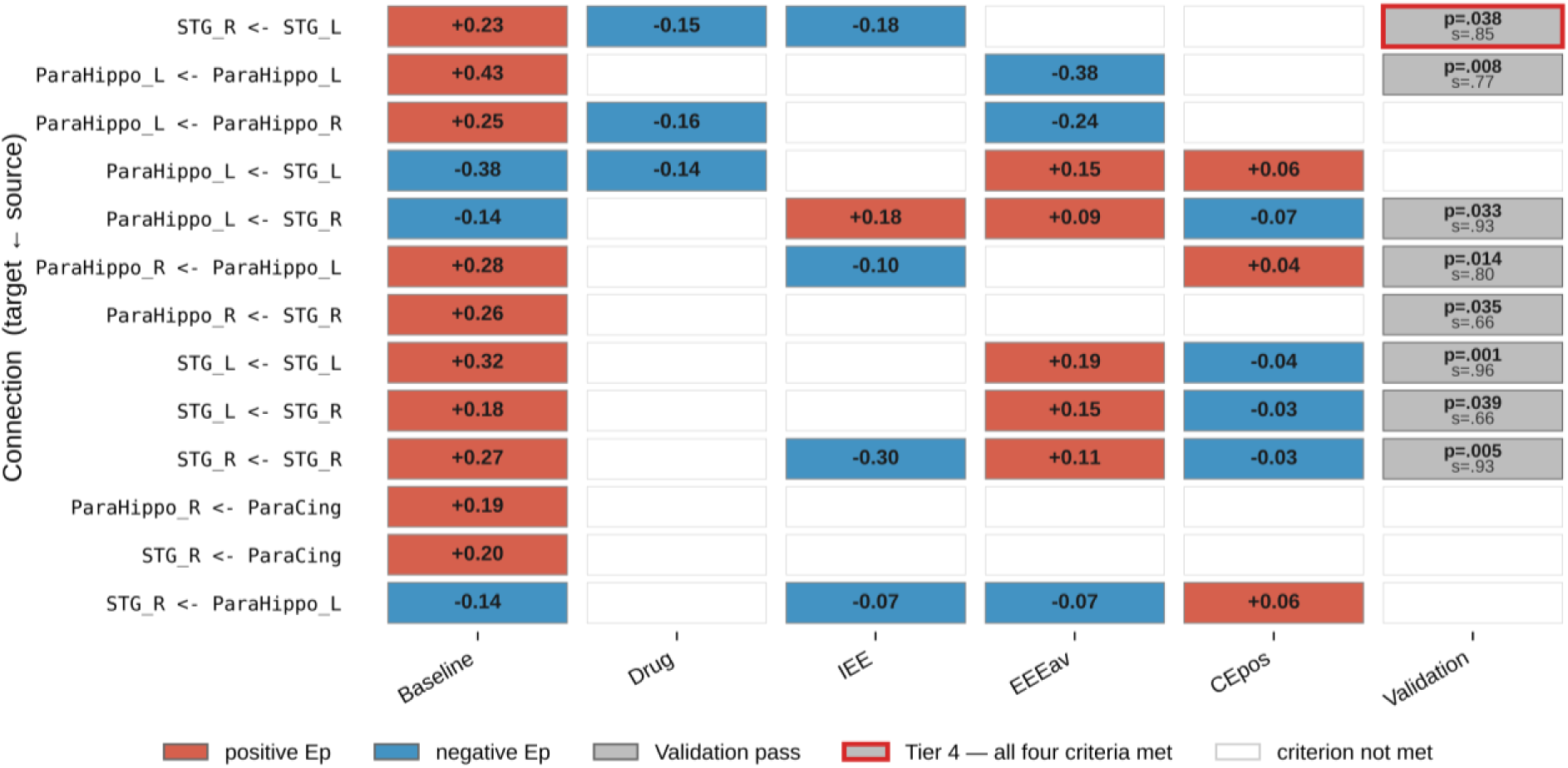
Neurosynth-derived DCM convergence framework. Each row is an effective-connectivity edge (target ← source) that reached Pp ≥ 0.99 (Bayesian local FDR ≤ 0.01) in at least one PEB design. Columns: Baseline (intercept of drug-effect PEB); Drug (drug regressor of drug-effect PEB); emotional arousal (IEE), empathic concern (EEEav), cognitive empathy (Cepos) (behavioral-association PEB regressors; a column fires if Pp ≥ 0.99 in either the PSIL or the PLA combined PEB); Validation (connection survived the PEB-level 1000-permutation null on |Ep| at p < .05 AND retained Pp ≥ 0.99 in ≥ 50% of 1000 subject bootstrap resamples in at least one behavioral sub-test; printed values are taken from the subscale condition test passing both criteria, selecting the smallest p when several pass). A red border on the Validation tile marks connections that met all three convergence criteria (Baseline, Drug, ≥ 1 behavioral sub-test) and additionally survived permutation and bootstrap validation. For self-connections (diagonal entries in EC matrices), Ep is a log-scaling factor on an intrinsically inhibitory term, so positive Ep (red) at baseline reflects stronger self-inhibition.

